# Genomic signatures of UV resistance evolution in *Escherichia coli* depend on the growth phase during exposure

**DOI:** 10.1101/2021.01.05.425512

**Authors:** S Selveshwari, Kasturi Lele, Sutirth Dey

## Abstract

Physiological states can determine the ability of organisms to handle stress. Does this mean that the same selection pressure will lead to different evolutionary outcomes, depending on the organisms’ physiological state? If yes, what will be the genomic signatures of such adaptation(s)? We used experimental evolution in *Escherichia coli* followed by whole-genome whole-population sequencing to investigate these questions. The sensitivity of *Escherichia coli* to ultraviolet (UV) radiation depends on the growth phase during which it experiences the radiation. We evolved replicate *E. coli* populations under two different conditions of UV exposures, namely exposure during the lag and the exponential growth phases. Initially, the UV sensitivity of the ancestor was greater during the exponential phase than the lag phase. However, at the end of 100 cycles of exposure, UV resistance evolved to similar extents in both treatments. Genome analysis showed that mutations in genes involved in DNA repair, cell membrane structure and RNA polymerase were common in both treatments. However, different functional groups were found mutated in populations experiencing lag and exponential UV treatment. In the former, genes involved in transcriptional and translational regulations and cellular transport were mutated, whereas the latter treatment showed mutations in genes involved in signal transduction and cell adhesion. Interestingly, the treatments showed no phenotypic differences in a number of novel environments. Taken together, these results suggest that selection pressures at different physiological stages can lead to differences in the genomic signatures of adaptation, which need not necessarily translate into observable phenotypic differences.

## 1. Introduction

Environmental stresses can strongly influence the growth and physiology of microbial cells (Aertsen & Michiels, 2004; Schimel *et al.*, 2007). At the same time, antecedent growth conditions and physiology of the cells can influence a microbe’s response to environmental stresses (Gilbert *et al.*, 1990; Lindqvist & Barmark, 2014; Lin & Kussell, 2016). For example, it has been shown that the response of bacteria to ionizing radiations are influenced by both the growth phases during which the exposure happens as well as the physiology of microbial cells. The typical batch culture involves bacterial growth through three distinct growth phases: lag (where cells adapt to the new growth conditions), exponential (characterized by rapid cell division) and stationary phase (where growth plateaus as a result of nutrient depletion) (Monod, 1949). Among the three growth phases, the fast dividing cells (in exponential phase) have been shown to be more resistant to ionizing radiations than growth arrested and non-dividing cells (in stationary phase) in *Moraxella spp*. (Keller & Maxcy, 1984), *Lactobacillus spp*. (Hastings *et al.*, 1986), *Halobacterium spp*. (Kottemann *et al.*, 2005; DeVeaux *et al.*, 2007), and yeast (Fabre, 1970). However, the converse has also been observed in species like *Escherichia coli* (Morton & Haynes, 1969; Dantur & Pizarro, 2004; Bucheli-Witschel *et al.*, 2010), *Cronobacter sakazakii* (Arroyo *et al.*, 2012), and *Salmonella typhimurium* (Child *et al.*, 2002). Although the effects of radiation on the exponential phase and the stationary phase have been explored in detail in the microbial literature, the same cannot be said about another important part of the microbial growth curve, namely the lag phase. Lag phase represents the adaptive period where the cells are metabolically active, but cell division has not begun (Rolfe *et al.*, 2012). Study of this phase of growth has a number of implications including food preservation (Sun, 2011), bacterial infections and antibiotic resistance (Frimodt-Møller *et al.*, 1983) as well as in maintaining laboratory cultures. Several environmental and physiological attributes that are unique to the lag phase (Bertrand, 2019 and the references therein) can influence the response to radiation stress (Gayán *et al.*, 2014). For example, microbes in lag phase express distinctive transcriptomic and proteomic profiles which include components essential for metabolism and growth (Rechinger *et al.*, 2000; Hornbæk *et al.*, 2004; Larsen *et al.*, 2006) as well as genes and proteins involved in repair of stasis induced macromolecular damages (Rolfe *et al.*, 2012). This increased expression of DNA repair machinery in the lag phase might influence the cells’ response to radiation. Consequently, the physiological state of the cell can influence its sensitivity to radiation both in the short term as well as over longer, evolutionary timescales.

Experimental evolution of resistance to ultraviolet radiation have been demonstrated in several microbial species including *Escherichia coli* (Ewing, 1995; Alcantara-Diaz *et al.*, 2004; Goldman & Travisano, 2011; Shibai *et al.*, 2017), *Salmonella typhimurium* (Davies & Sinskey, 1973), *Bacillus subtilis* (Wassmann *et al.*, 2010), *Pseudomonas cichorii* (Weigand & Sundin, 2009), and T7 bacteriophage (Tom *et al.*, 2018). Although all these studies found that repeated exposures to UV resulted in increased ability to survive UV stress, there were several differences in the correlated responses to selection. For example, the increase in UV resistance was accompanied by increased cell size (Goldman & Travisano, 2011), increased tolerance to osmotic, oxidative and desiccation stress (Wassmann *et al.*, 2010), and the appearance and maintenance of rare colony morphologies due to increased rate of UV induced mutations (Weigand & Sundin, 2009). These studies on evolution of radiation resistance looked at the effects of exposure either during the lag phase (Ewing, 1995; Weigand & Sundin, 2009; Goldman & Travisano, 2011; Shibai *et al.*, 2017) or the stationary phase (Davies & Sinskey, 1973; Alcantara-Diaz *et al.*, 2004; Wassmann *et al.*, 2010). Consequently, there is little understanding of the evolutionary effects of exposure during the exponential phase. At the same time, these existing studies have primarily focused on the evolution of resistant phenotypes and the correlated phenotypic changes (but see Shibai *et al.*, 2017 and Tom *et al.*, 2018). The genomics of radiation resistance has been studied in detail in radio-resistant species such as Deinococcus spp. (White *et al.*, 1999; Blasius *et al.*, 2008), Rhodobacter spp. (Perez *et al.*, 2017). However, there is relatively less work on this aspect in the context of non-radio-resistant species (although see Bruckbauer *et al.*, 2019). It is not obvious that the insights gained about the genomic correlates of exposure to UV in the above radio-resistant species, would be applicable to other microbes. Thus, in order to unearth the putative mechanisms underlying evolution of radiation resistance, it is important to extend the genetic studies conducted in the radio-resistant species to other microbes.

In this study, we attempt to address some of these lacunae in our understanding of the evolution of radio-resistance, particularly ultraviolet radiation. First, we observe that our ancestral population of *Escherichia coli* are more sensitive to UV in the exponential phase than in lag phase. To study the evolutionary response to UV radiation in these two phases, we subjected two sets of replicate populations to 100 rounds of UV exposure and growth in the lag and the exponential phases. Both treatments showed significant reduction in sensitivity to UV compared to control populations. However, growth phase differences in UV sensitivity were no longer observed. To investigate the genomic correlates of evolution of UV resistance, the UV-treated and control populations were further subjected to whole-genome whole-population sequencing. Genes associated with DNA repair pathways, RNA polymerase and cell membrane structure were commonly mutated in both UV-treated populations. Additionally, mutations in the populations exposed to UV in the lag phase were grouped in genes involved in transcription / translation regulation and cellular transport. On the other hand, only the populations that faced UV in the exponential phase contained mutations in signal transduction and cell adhesion. The genes and pathways that were mutated during UV resistance evolution may also have led to correlated changes in growth and survival in novel environments such as antibiotics, heavy metal salts and minimal media with a single carbon source. Contrary to our expectation, these differences in the genome did not translate to phenotypic differences between lag and exponential treatments in either the selection or the novel environments.

## 2. Materials and Methods

We used an *Escherichia coli* K12 MG1655 strain in which the lacY gene had been replaced with a Kanamycin resistance gene. Colonies of this bacterium are white coloured on MacConkey’s agar as opposed to the red coloured colonies produced by other *Escherichia coli*. All cultures were maintained at 37°C and 150 RPM throughout the selection and assays, except where stated otherwise. An overview of the experimental design is given in a supplementary figure (Figure S1).

### 2.1 Experimental evolution

Six populations were initiated from six independent *E. coli* colonies picked from a nutrient agar streaking of the ancestral *E. coli* strain. These six are henceforth referred to as the ancestor populations. From each ancestor, we derived three replicate populations and assigned them randomly to one of the three selection regimes, namely lag, exponential and control. Both lag and exponential treatments were subjected to UV during the respective growth phases while the control was devoid of UV exposure. The populations were exposed to UV in a custom-built dark chamber where the only source of light was an 8W UV-C tube-light (Philips TUV 8W) with an emission peak at 254 nm. The lamp was placed at a height of 18 inch from a platform shaker, which results in a constant irradiance of 100.5 μW/cm^2^. At the beginning of the experiment, the duration of exposure was 15 seconds, which resulted in 2 – 3 log_10_ reduction (100 – 1000 fold) in colony forming units (CFUs); see section 2.2 for more details regarding estimation of log_10_ reduction. To allow sufficient time for the populations to adapt, the duration of UV exposure was increased gradually i.e., once every 5 days. For the full trajectory of exposure duration throughout selection, from day 0 – 100, see supplementary table (Table S2). By the end of 100 days of evolution, the exposure time was 370 seconds, i.e. an increase of ~25 times.

All cultures were grown in 2ml nutrient broth with kanamycin (NB^Kan^) in six-well tissue culture plates (see supplementary Table S3 for composition). The use of six-well plates allowed for a larger surface area to volume ratio for better UV penetration. Growth of all populations reached a plateau within 24 hours. Every 24 hours, 20 μl of the grown culture was sub-cultured into fresh NB^Kan^. Control populations (C) were sub-cultured in NB^Kan^ with no UV exposures whereas, populations designated as Lag (L) treatment were exposed to UV during the lag phase, immediately after subculture (0h). Exponential (E) treatment populations were exposed to UV during the exponential phase by monitoring their growth/OD_600_ in a plate reader (Synergy HT, BIOTEK Winooski, VT, USA) at 20-minute intervals. Based on pilot experiments, the exponential phase was deemed to have been reached when the OD_600_ of the culture was greater than 0.11 and the difference between two consecutive OD measurements were greater than 5% for the first time, i.e. OD_t+1_ >1.05 × OD_t_, where the subscript t and t+1 refer to two successive time points. Populations in the exponential treatment were exposed to UV only when both the conditions were met. The 6-well plates containing the cultures were placed on the shaker platform directly under the UV lamp without the lid. The cultures were shaken at room temperature and 150 RPM during exposure. Since bacteria possess the photoreactivation pathway which can induce error-free reversal of UV induced damages and reduce the efficiency of UV radiation (Thoma, 1999; Sinha & Hader, 2002), the cultures were maintained in darkness throughout the process of selection, except during sub-culturing. The UV induced mortality makes it impossible to accurately estimate the dilution ratio and consequently the number of generations in the UV-treated populations. However, at the end of 100 days, the UV-treated populations would have experienced more generations, on average, than control populations which underwent ~667 generations of evolution (6.67 doublings/transfer x 100 days). Every 5 days, 15% glycerol stocks (300 μl of 50% glycerol + 700 μl culture) of all populations was prepared and stored at −80°C. Due to logistic reasons, selection was interrupted once at day 60. Selection was reinitiated by reviving 10μl glycerol stocks of all populations (UV-treated and controls) in 90 μl NB^Kan^ in a 96 well plate for 12 hours. 20 μl of this revived culture was inoculated in 2ml NB^Kan^ to initiate day 61. Glycerol stocks prepared at the end of 100 days, were used for all assays.

### 2.2 Measuring UV sensitivity

Sensitivity to UV induced mortality was measured as the log of change in number of colony forming units (CFUs) before and after UV exposure (log_10_(CFUs before exposure/CFUs after UV exposure)) (Koivunen & Heinonen-Tanski, 2005).The populations to be assayed were revived by inoculating 5 μl of the corresponding glycerol stocks in 2ml NB^Kan^ and incubating them overnight at 37°C. Assays were conducted in six well plates with 20μl of the revived culture inoculated in 2ml NB^Kan^. UV sensitivities of all populations were assayed in both lag and exponential phase where the UV exposures were carried out in a manner similar to the selection procedure. CFU counts before and after exposure were determined by serially diluting the cultures and plating 100 μl of the appropriate dilutions (to obtain countable number of colonies) on 2% nutrient agar containing kanamycin (NA^Kan^). The serial dilution and plating after UV exposure was carried out in a dark room illuminated by red light to prevent photo-reactivation. The NA^Kan^ plates were incubated in darkness at 37°C, overnight and the number of colonies were counted manually and multiplied by the dilution factor to obtain the number of CFUs.

To measure the UV sensitivities of the ancestors, all six ancestral populations were assayed at both lag and exponential phase and at four exposure durations: 15, 60, 200, and 370 seconds. After 100 rounds of selection for UV resistance, we measured the UV sensitivity of the evolved populations L, E, and C along with the ancestors. Log_10_ reduction in CFUs during both lag and exponential growth phases was measured at exposure duration of 370 seconds. Two independent measurement replicates of the UV sensitivities of the evolved populations were obtained by conducting the entire assay from revival to CFU counts, twice. The relative change in UV sensitivity due to selection was obtained by scaling the sensitivities of the evolved populations by the sensitivity of the ancestors in the corresponding growth phases.

### 2.3 Whole-genome sequencing and analysis

To understand the genomics of repeated exposures to UV and selection for UV resistance, we randomly chose two replicates (rep 1 and 3) of the evolved populations (L, E, C), and their corresponding ancestors, and subjected them to whole-genome whole-population sequencing. 5 μl of the corresponding glycerol stocks were inoculated in 4ml NB^Kan^ and incubated overnight at 37°C. 3ml of the revived cultures were centrifuged down, washed twice in phosphate buffer saline (PBS), air dried and shipped for genome sequencing to a commercial service-provider. They performed paired-end whole-genome sequencing on NextSeq500 platform (Illumina, USA) at ~100X (range: 98.8X – 139.2X) coverage and 150bp read length. The service-provider provided us with high quality reads after removing adaptor sequence, ambiguous reads (reads with unknown nucleotide “N” of more than 5%), and low-quality sequences (reads with more than 10% having a phred score < 20) using Trimmomatic v0.38. We then used *Breseq* version 0.33.2 pipeline (Deatherage & Barrick, 2014) at default parameters for sequence alignment and variant calling. First, mutations in the ancestral genome were identified (in consensus mode) by aligning it to the reference genome of *Escherichia coli* MG1655 (Genbank accession: NC_000913.3). Breseq’s gdtools package was used to incorporate the predicted mutations and update the ancestral genome. This updated ancestral genome sequence was then used as reference for alignment and variant calling (in polymorphism mode) in the evolved populations. Following previous studies (Bailey *et al.*, 2015; Sandberg *et al.*, 2017; Santos-Lopez *et al.*, 2019), the list of predicted mutations were further curated by removing all variants present at frequencies less than 10%. We also aligned the ancestral sequence in the polymorphism mode. To limit our analysis to the genomic changes that evolved *de novo* in response to selection, all polymorphic variants common between the ancestral and evolved populations were also removed from the dataset. This curated list of mutations was used for computing the number of SNPs and indels, SNPs in coding vs intergenic regions, synonymous vs nonsynonymous SNPs, as well as the mutational spectrum of all mutations. We estimated the dN/dS ratio as the number of non-synonymous mutations per non-synonymous site (dN) to the number of synonymous mutations per synonymous sites (dS). The total number of non-synonymous and synonymous sites in the genome was estimated by using Breseq’s gdtools package. Note that the relationship between dN/dS ratio and the selection coefficient can be an issue when comparing two populations sharing a number of fixed mutations. However, this issue is avoided in our study as we compare evolved populations with their ancestors (Chen & Zhang, 2020). Functional annotations and enrichment analysis of the genes were carried out on DAVID v.6.8 (Huang *et al.*, 2009; Sherman & Lempicki, 2009) a web-based bioinformatics application. The list of mutated genes was classified into functional groups (based on UniProtKB keywords) followed by manual curation.

### 2.4 Measuring fitnesses in novel environment

We observed that the UV-treated populations had accumulated numerous mutations across the genome. These UV induced mutations could possibly affect the fitness of the populations under environmental conditions not faced during selection (see Discussion for details). Therefore, after 100 days of selection for UV resistance, we assayed the fitness of the evolved populations (L, E, C) as change in minimum inhibitory concentration (MIC) and growth rate in a suite of environments.

#### Measuring minimum inhibitory concentrations

MICs of the evolved populations were measured in four antibiotic environments and three heavy metal environments. Antibiotics (ampicillin, chloramphenicol, nalidixic acid and rifampicin) from four different classes were chosen for their different mechanisms of action (Kohanski *et al.*, 2010). Similarly, the heavy metal salts (cobalt chloride, copper sulphate and nickel chloride) were chosen for the different ways in which they disrupt metabolic processes (Dupont *et al.*, 2011; Macomber & Hausinger, 2011; Majtan *et al.*, 2011). To the best of our knowledge, evolution of resistance to UV has no known association to fitness in these environments.

Populations were grown in increasing concentrations of the assay environment (antibiotics or heavy metals), and the minimum concentration at which no visible growth was obtained was taken as MIC. The evolved populations and their corresponding ancestor were revived as mentioned before. A gradient of the assay environments was prepared by serial two-fold dilutions in 96 well plates. The revived cultures were inoculated in each concentration of the assay environment at a dilution of 1/1000 in triplicates. After 48 hours of growth, each plate was scored for absence of growth either visually (for antibiotic environments) or when OD_600_ < 0.2 (for heavy metals). A concentration was considered the MIC of the population only if at least two out of the three replicates showed no growth. The entire assay from revival to MIC determination was repeated twice and served as measurement replicates. MIC of all the evolved populations were scaled by the MIC of their corresponding ancestor to obtain the change in MIC due to selection.

#### Measuring growth rate

Growth rates of the evolved populations were assayed under two kinds of conditions: a) in nutrient broth, and b) in M9 minimal media containing a single carbon source (for composition, see supplementary Table S4). Growth rate was measured in five different carbon sources: fructose, glucose, glycerol, mannose and thymidine, all of which feed into the glycolysis pathway via different intermediates (Voet & Voet, 2011). Mutations in the intermediate enzymatic steps can likely alter the efficiency of metabolism of the carbon sources and consequently growth. The revived evolved and ancestral populations were inoculated in 200 μl of the assay environment (i.e. M9 media containing one of the five carbon sources) at a dilution of 1/1000 in 96 well plates. These cultures were subjected to automated growth measurements using a plate reader (Synergy HT, BIOTEK Winooski, VT, USA). OD_600_ was measured every 20 minutes for 24 hours at 37°C and slow continuous shaking, at 17 cycles per second. Following previous studies (Karve *et al.*, 2015; Sprouffske *et al.*, 2018; Chavhan *et al.*, 2019; Rodríguez-Rojas *et al.*, 2020), we computed the growth rate as the maximum slope of the curve over a moving window of 10 readings. Two measurement replicates were obtained by repeating the entire assay twice. The growth rates of the evolved populations were scaled by the ancestral growth rate to obtain the relative change in growth rate due to selection.

### 2.5 Statistical analysis

Ancestral UV sensitivities were compared using a three-way mixed model ANOVA with randomized complete block design (RCBD) (Rohlf & Sokal, 1980). Here, we used growth phase (lag and exponential) and exposure duration (15s, 60s, 200s, and 370s) as fixed factors crossed with each other and replicate populations (6 populations) as an independent random factor (neither crossed nor nested in other factors). Cohen’s *d* (Cohen, 2013) was computed to assess the effect sizes of the differences between the two growth phases. The effects sizes were interpreted as small, medium and large for 0.2 < *d* < 0.5, 0.5 < *d* < 0.8 and *d* > 0.8, respectively (Sullivan & Feinn, 2012). UV sensitivities of the evolved populations after 100 days of selection were compared in a three-way mixed model ANOVA with selection (lag, exponential and control) and assay environment (lag and exponential) as fixed factors and replicate populations (6 populations) as a random factor in a full factorial design. Fitness in novel environments were analyzed as separate two-way mixed model ANOVAs for each fitness measure and environment. Selection (lag, exponential and control) as fixed factors and replicate populations (6 populations) as a random factor were analyzed in a full factorial design. Correction for inflation of family-wise error rate was done for the two fitness measurements, independently, using the Holm–Šidák procedure (Abdi, 2010).

We compared the dN/dS ratio of the evolved populations to the expected ratio (0.754, ratio of possible synonymous sites to nonsynonymous sites in the genome computed using Breseq) using a binomial test on R 3.6.1.

All the ANOVAs were performed on STATISTICA v7.0 (Statsoft Inc.). Cohen’s *d* statistics were estimated using the freeware Effect Size Generator v2.3.0 (Devilly, 2004).

## 3. Results

### 3.1 Ancestral UV sensitivity in lag and exponential growth phases

We first measured the UV sensitivity of the ancestral *Escherichia coli* during the lag and exponential phases at four exposure durations. We found a significant interaction between growth phase and exposure duration (*F*_3,35_ = 3.34, *P* = 0.03). The exponential phase sensitivity was significantly larger than the lag phase sensitivity in all but one comparison in Tukey’s post hoc analysis (Fig. 1; 60s: p=0.0005; Cohen’s *d* = 2.69 (large), 200s: p=0.002; Cohen’s *d* = 2.93 (large), 370s: p=0.0001; Cohen’s *d* = 4.4 (large)). At 15 seconds of exposure, the difference between lag and exponential exposure was marginally non-significant but with a large effect size (Fig. 1; 15s: p=0.065; Cohen’s *d* = 2.09 (large)). Additionally, the six ancestral populations show no significant differences in their UV sensitivity (*F*_5,35_ = 0.36, *P* = 0.87). Together, this shows that our ancestral strain of *E.coli* was more sensitive to UV during exponential phase of growth. However, as the interaction of growth phase and exposure was significant, we refrain from interpreting the main effects.

**Figure 1.**
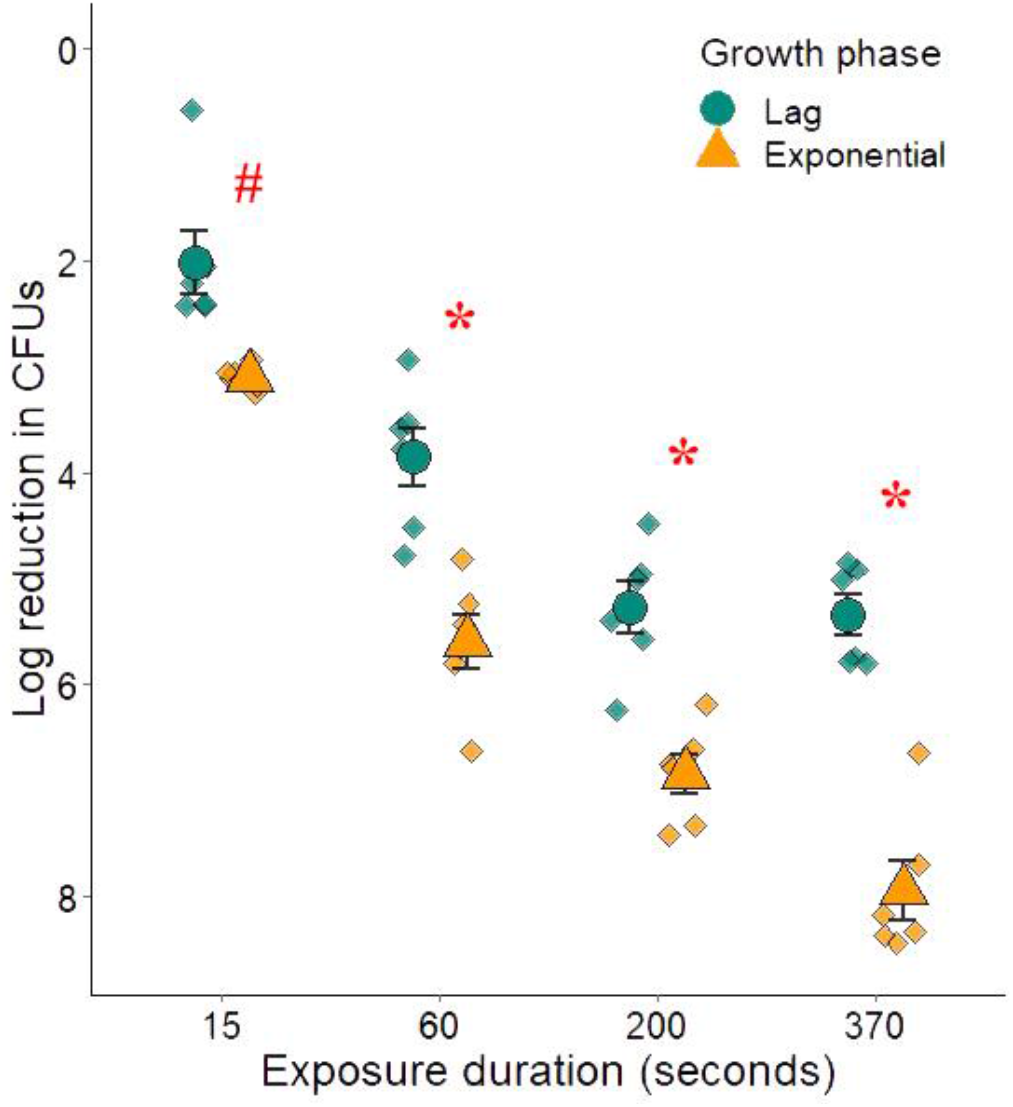
UV induced log_10_ reduction in CFUs during lag and exponential growth phases. UV sensitivity of the ancestral *E.coli* was measured at four exposure durations (15, 60, 200, and 370 seconds) and two growth phases. Mean sensitivity of the six ancestral populations are plotted as circles (lag phase) and triangles (exponential phase). Whiskers represent ±SE. The scatter of the six ancestral value is represented by diamonds (♦). * denote p value < 0.05 in Tukey’s post-hoc analysis at that exposure value and # denotes p = 0.065

### 3.2 UV sensitivity of evolved populations

UV sensitivity of the evolved populations in the two growth phases were compared after scaling them by the corresponding ancestral values. At the end of 100 rounds of UV exposure and selection, the populations significantly differed in their relative UV sensitivities (*F*_2,36_ = 249.72, *P* = 2.91E-09). Tukey’s post hoc analysis showed that both lag (L) and exponential (E) populations had evolved significantly higher resistance compared to control (C) populations but were not different from each other (Fig. 2; L and C *P* = 0.0001; E and C *P* = 0.0001 and L and E *P* = 0.13). Interestingly, the interaction between selection lines and assay environments was also not significant (*F*_2,36_ = 3.82, *P* = 0.059). This also suggests that the populations had evolved similar extents of resistance in both growth phases irrespective of the selection environment.

**Figure 2.**
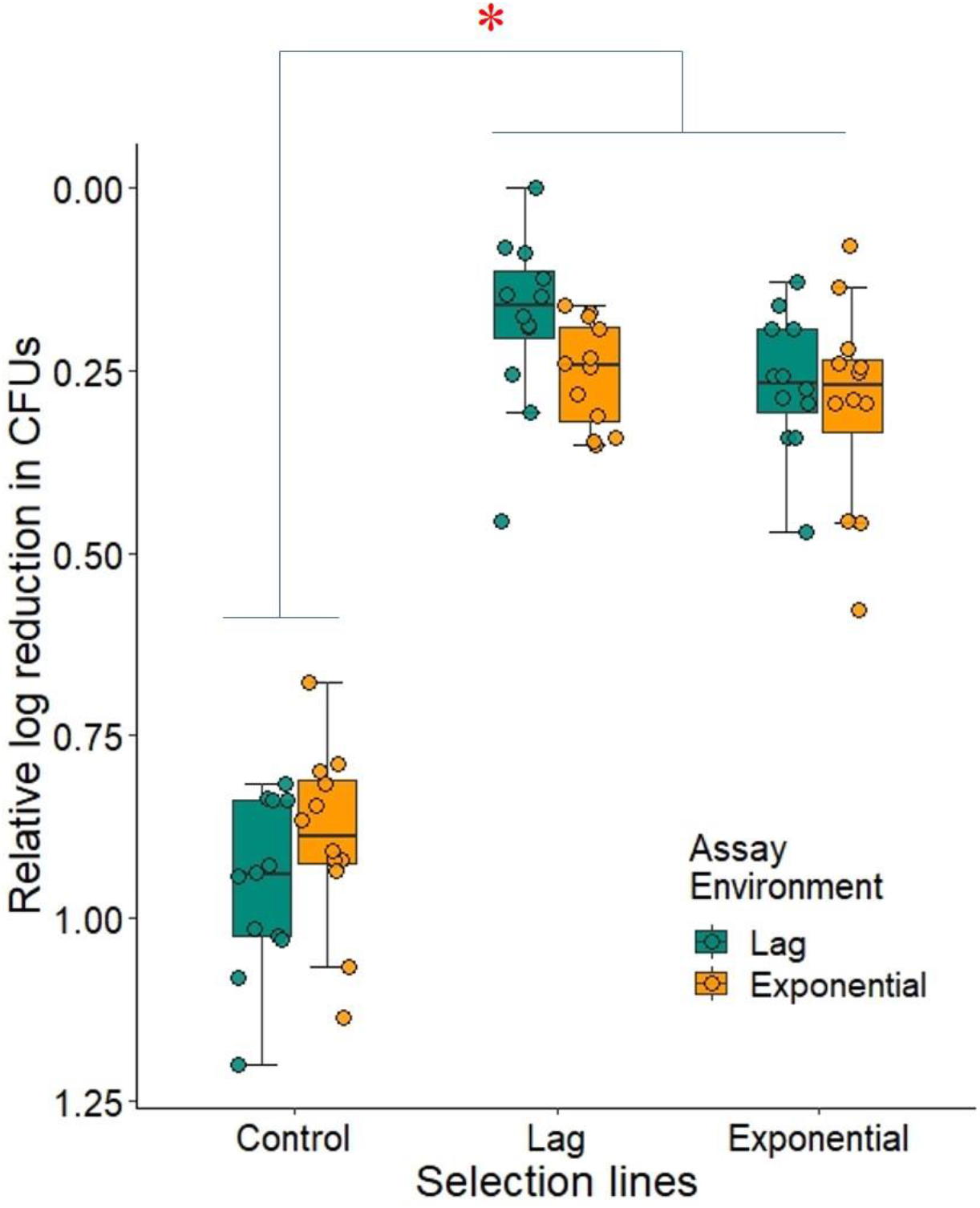
UV sensitivity of evolved populations relative to the ancestor’s sensitivity. Decline in the relative log_10_ reduction (sensitivity to UV) in lag and exponential populations signifies evolution of resistance. Each box plot represents data from 12 values i.e., 6 replicate population assayed twice. Solid lines represent median, upper whisker denotes the largest data point lesser than or equal to 1.5 * IQR (inter-quartile range) and similarly for the lower whisker. * denotes that control population has significantly (p < 0.05) lower resistance to UV than both lag and exponential populations in Tukey’s post-hoc analysis.

### 3.3 Genomics of UV resistance evolution

To investigate the genomic changes accumulated during selection for UV resistance, we carried out whole-genome whole-population sequencing of two replicate populations from each selection regime (L, E, C) and their corresponding ancestors. Both our ancestral populations had nearly identical sequences. However, our lab strain differed from *E.coli* K-12 reference genome (NC_000913.3) at several genome locations. Therefore, we used the assembled sequences of the corresponding ancestor as reference for identifying mutations in the evolved populations. Exposure to UV in both growth phases resulted in a marked increase in the total number of mutations. The two replicates of lag (193 and 317) and exponential (60, 319) treatments had much greater number of single nucleotide polymorphisms (SNPs) compared to the control populations (8, 13). A histogram showing the distribution of mutation frequencies of all SNP in the four UV-treated populations is given in the supplementary figure S8. Very few indels (insertions and deletions) were identified in all three evolved populations (2, 2 in lag, 4,12 in exponential and 3,7 in control). Some of these indels, 0 (L1), 1 (L3, E1, E3), 2 (C1) and 5 (C3), were found to be mediated by IS2 insertional elements.

We also found that the mutational spectra of both lag and exponential treatments were transition-biased as GC→AT transitions accounted for at least 50% of all mutations, a confirmed signature of exposure to UV (Fig. 3) (Griffiths *et al.*, 2005; Brash, 2015). In spite of the bias, all six mutations types were represented in both UV treatments. Mutations in the control populations were confined to three or four types only (see supplementary table S5).

**Figure 3:**
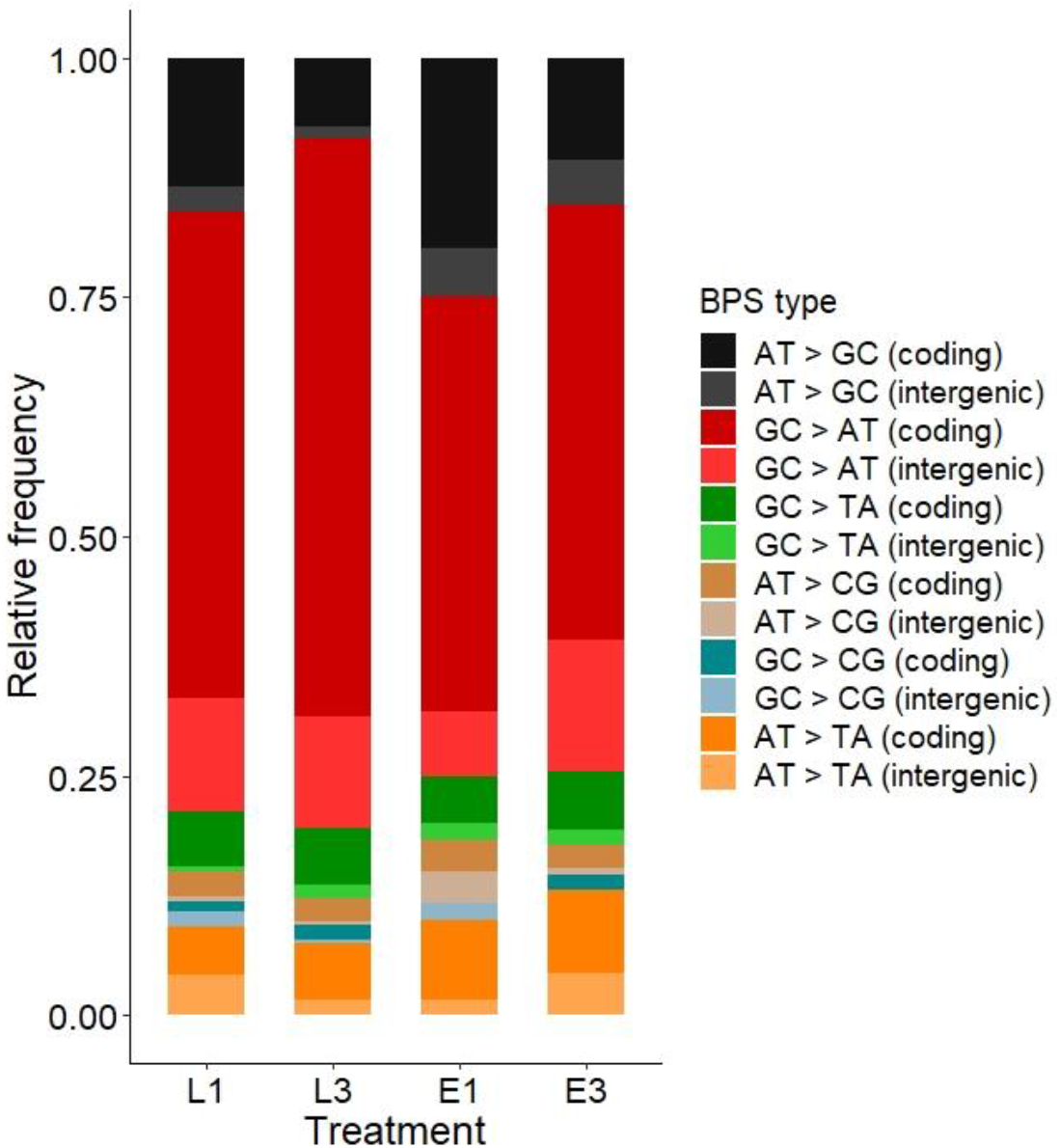
Mutational spectrum of SNPs accumulated during lag and exponential UV exposure. Colours in the stacked bar represent the relative frequency of each of the six type of base-pair substitutions (BPS), in the coding and intergenic regions‥

We notice that 75 – 80% of all SNPs in the UV treatments were found in the coding regions which typically makes up about 88% of the *Escherichia coli* genome (Rogozin *et al.*, 2002). We estimated the dN/dS ratio (ratio of non-synonymous mutations per non-synonymous sites to synonymous mutations per synonymous sites) to infer the selection pressure that led to the genomic changes. Table 1 summarizes the dN/dS ratios in the evolved populations. The ratios were statistically significant in both the lag treatments (L1, L3) and one of the exponential treatment (E3). In all three cases, the ratios were less than unity, indicating that these populations were primarily subjected to purifying selection (Yang & Bielawski, 2000).

**Table 1:**
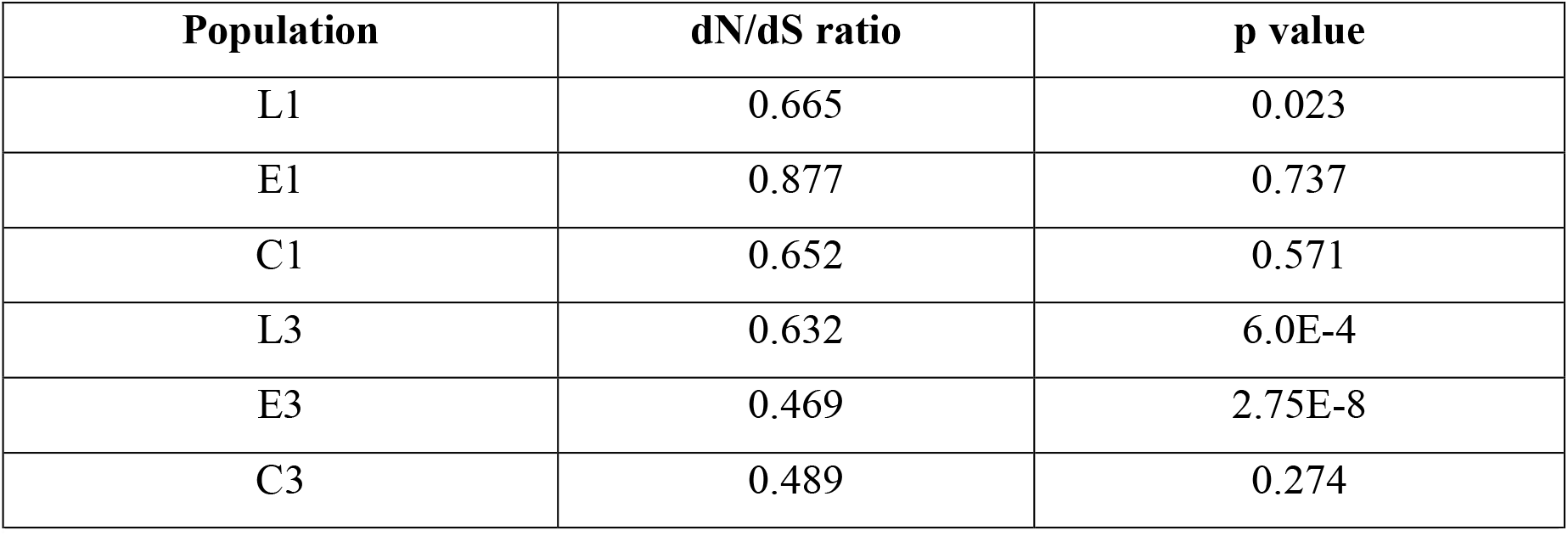
Estimates of non-synonymous mutation to synonymous mutation ratio in the evolved populations. p value denotes the binomial probability that the observed dN/dS ratio differs from the expected ratio (0.754), ratio of possible synonymous sites to nonsynonymous sites in the genome)

Next, we investigated the genetics of UV resistance evolution by considering only the nonsynonymous mutations in the UV-treated populations. To begin with, the genes that were mutated differed considerably between lag and exponential treatments (see supplementary file S9 for full list of non-synonymous mutations). There were only two genes (*recA* and *mepS*) that had mutated in both treatments but not in the control populations. Our ancestral populations had a single mutation in *recA* (G161D) with respect to the reference MG1655 strain. Reversion of this mutation had fixed in all UV-treated populations but not the controls. We note that this reversion in *recA* (D161G) is the only consistent mutation where all treatments have the same amino acid change. In addition to this, one other mutation (P314L) in *recA* was present at low frequency in L3. *recA* is involved in DNA recombination and repair (Smith *et al.*, 1987; Schlesinger, 2007) and also plays an integral role in the mediation of SOS response (Markham *et al.*, 1985; Maslowska *et al.*, 2019). The second commonly mutated gene was *mepS* with four different mutations: Y82I (in E1), R83C (in L1), P111L (in L3, E3), and T114I (in L3). *mepS* is known to be involved in peptidoglycan biosynthesis during cell growth (Singh *et al.*, 2012). Populations L1 and E3 had mutations in recJ and L3 had mutation in recQ, both components of the recFOR recombination pathway. Similarly, RNA polymerase genes rpoC was mutated in L1 and E3 population, while mutation in rpoB was observed n L3.

Besides these commonly mutated genes, we also identified 9 genes that were mutated in both the replicates of lag treatment. Of these, a functional cluster of 5 genes was identified. *crp*, *deoR*, *fadR*, *hfq*, and *lexA* are known to be involved in DNA-binding, transcription and translational regulation. Additionally, *lexA* in association with *recA,* is known to mediate the SOS response. The remaining four genes show no obvious clustering (see Table 2 for full functional annotations). Similarly, in the populations exposed to UV during the exponential phase, we identified four genes that were mutated in both replicates. However, these genes too showed no obvious clustering of function (Table 2).

**Table 2:** List of genes consistently mutated in both replicates of lag and exponential treatments. The genes along with their functional annotations are listed.

| **Lag treatment** |  |
| --- | --- |
| **Gene** | **Functional annotation** |
| *crp*<br>*deoR*<br>*fadR*<br>*hfq*<br>*lexA* | DNA-binding proteins, Repressors, Transcription and translational regulation |
| *lexA (recA)* | SOS response |
| *adiA* | Arginine catabolic processes |
| *fdoG* | Cellular respiration |
| *frlD* | Carbohydrate phosphorylation |
| *ycbK* | Uncharacterised protein |
| **Exponential treatment** |  |
| **Gene** | **Functional annotation** |
| *fimA* | Cell adhesion |
| *phoR* | Response to phosphate starvation and signal transduction |
| *ygfS* | Oxidation-reduction process |
| *yoaE* | Membrane component |

Using DAVID v.6.8, a web-based bioinformatics application, we carried out functional annotation and enrichment analysis of the list of non-consistent mutations. Only about half the list of genes fell into clusters of groups having similar functions (Fig. 4). This could have resulted in the observed lack of significance in enrichment analysis (supplementary Table S6). However, functional clusters comprising genes involved in or a part of cell membrane, were represented in three out of four populations (L1, L3, and E3). This, in addition to the fact that mutations of *mepS* were consistent across treatments, can suggest that cell membrane modification may play an important role in evolution of UV resistance. A second major group, that was found to be common between the two replicates of lag treatment, consisted of genes involved in transmembrane transport. Aside from these, mutations in lag treatment were grouped by genes involving in ligase activity, acetylation, tRNA-binding, nucleotide binding, and exopolysaccharide synthesis (Figure 4 and supplementary Table S6). Similarly, mutations in exponential treatments were grouped by genes involved in metal-ion binding, nucleotide binding, pyridoxal phosphate binding, kinase, and serine esterase. These differences imply that selection for UV resistance at the two growth phases can result in differences in how and where mutations accumulate in the genome.

**Figure 4:**
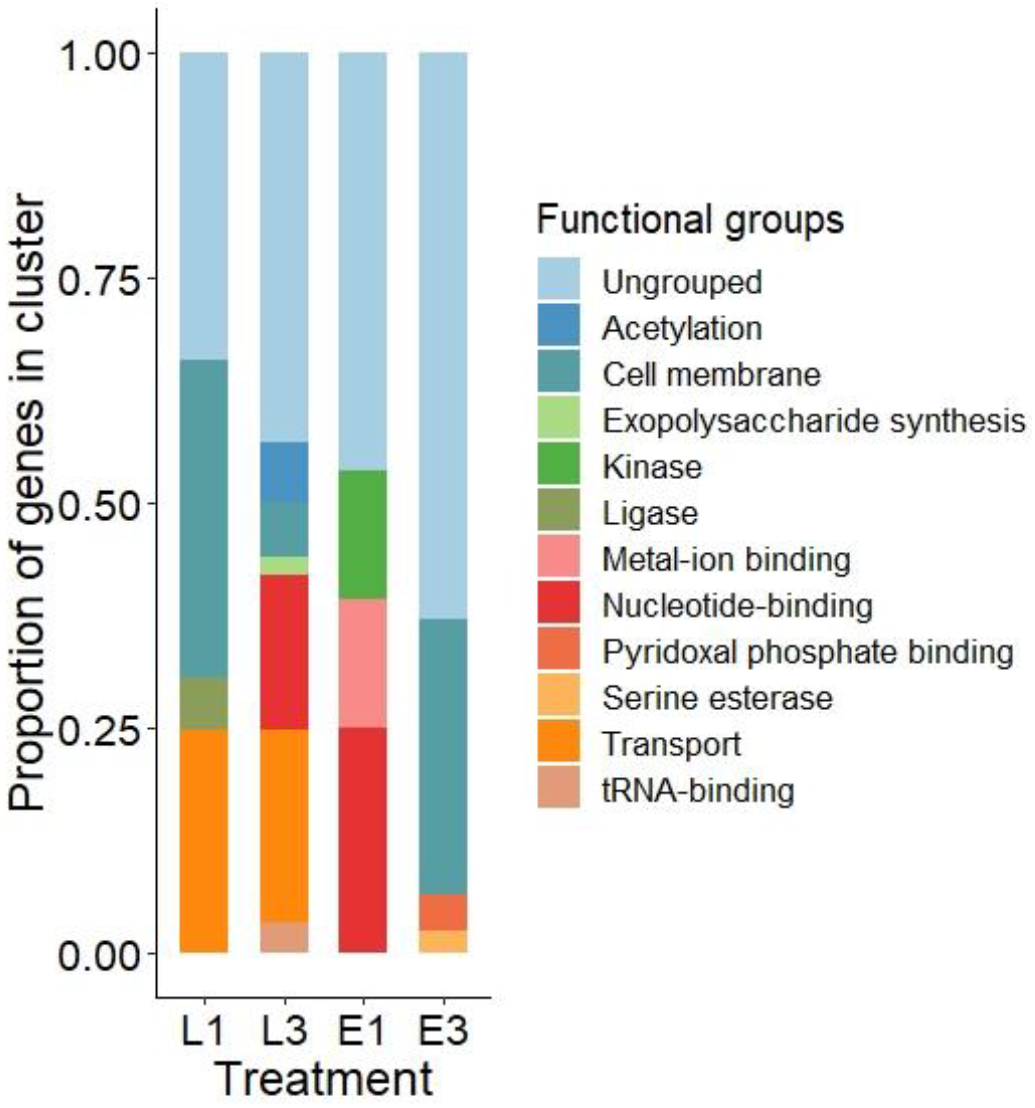
Functional grouping of non-consistent mutations. The functional categorizations are based on uniprot keywords and GO terms. Height of each stack represents the proportion of genes in that particular functional group. Functional annotation was done using DAVID v.6.8 followed by manual curation. See supplementary material Table S6 for list of genes in each group.

To summarize, repeated exposure to UV resulted the accumulation of a large number of mutations despite the effects of purifying selection. While genes such as *recA* and *mepS* were commonly mutated in all UV-treated populations, genes of different functional classes were mutated in lag and exponential treatments. This led us to the investigation of how these mutations affected the fitness of the populations under other environmental conditions.

### 3.4 Fitness in novel environment

Fitness of the evolved populations in novel environments was measured in two ways. Growth rate was measured in nutrient broth (NB^Kan^) and M9 minimal media in the presence of 5 different carbon sources: fructose (Fru), glucose (Glu), glycerol (Gly), mannose (Man) and thymidine (Thy). After correction for inflation of family-wise error rate (Holm–Šidák’s correction) only NB showed a significant effect of selection (Fig. 5A: *F*_2,18_ = 17.04, *P* = 0.0036). Tukey’s post-hoc analysis show that both lag and exponential treatments have increased growth rate in NB^Kan^ compared to control populations but did not differ w.r.t each other. We also measured the resistance of evolved population in stress environments. Resistance was measured as change in MIC in 4 antibiotic environments: ampicillin (Amp), chloramphenicol (Chl), nalidixic acid (Nal), and rifampicin (Rif) and 3 heavy metal environments: cobalt chloride (Co), copper sulphate (Cu), and nickel chloride (Ni). After correction for multiple testing (Holm–Šidák’s correction), main effect of selection was significant in chloramphenicol (Fig. 5B: *F*_2,18_ = 16.76, *P* = 0.0037), nalidixic acid (*F*_2,18_ = 70.33, *P* = 9.02E-06), rifampicin (*F*_2,18_ = 16.37, *P* = 0.0035), and cobalt chloride (*F*_2,18_ = 7.22, *P* = 0.045). In all four cases, Tukey’s post hoc tests suggested that lag and exponential treatments had increased MIC w.r.t control populations. Only in the case of chloramphenicol, there was a difference between lag and exponential populations: lag had higher MIC than exponential treatment (*P* = 0.046 in Tukey’s post hoc analysis). To further explore the difference in resistance between lag and exponential treatments, we also measured their growth rate at sub-MIC concentration (0.5 μg/ml) of chloramphenicol. The main effect of selection was significant (Fig. 6: *F*_2,18_ = 49.67, *P* = 6.4E-06) and both lag and exponential treatments had higher growth rate than control populations. But interestingly, populations in the exponential treatment had higher growth rate than those in the lag treatment (*P* = 0.0005 in Tukey’s post hoc analysis).

**Figure 5:**
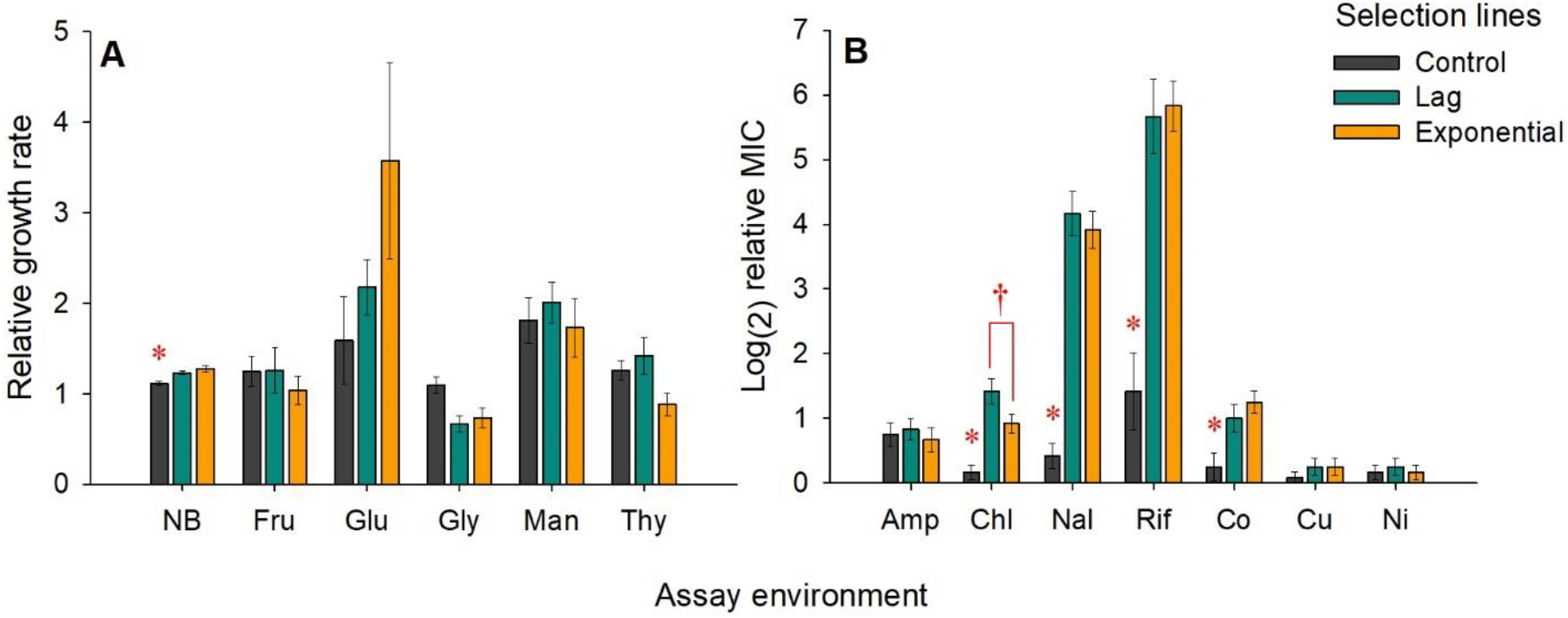
Relative fitness of the evolved populations in novel environment. Fitness is measured as (A) relative growth rate and (B) resistance, measured as minimum inhibitory concentration (MIC). Growth rate of the UV-treatedand control populations were assayed in nutrient broth (NB), M9 minimal media containing fructose (Fru), glucose (Glu), glycerol (Gly), mannose (Man) and thymidine (Thy). Resistance of all three evolved populations were measured in ampicillin (Amp), chloramphenicol (Chl), nalidixic acid (Nal), rifampicin (Rif), cobalt chloride (Co), copper sulphate (Cu), and nickel chloride (Ni). Fitnesses of the evolved populations were scaled by the ancestral fitness. Bars represents mean of 12 values i.e., 6 replicate populations assayed twice. Whiskers represent ±SE. * denote control is significantly (p value < 0.05) less than lag and exponential in Tukey’s post-hoc analysis. † denotes significant differences between lag and exponential treatments.

**Figure 6:**
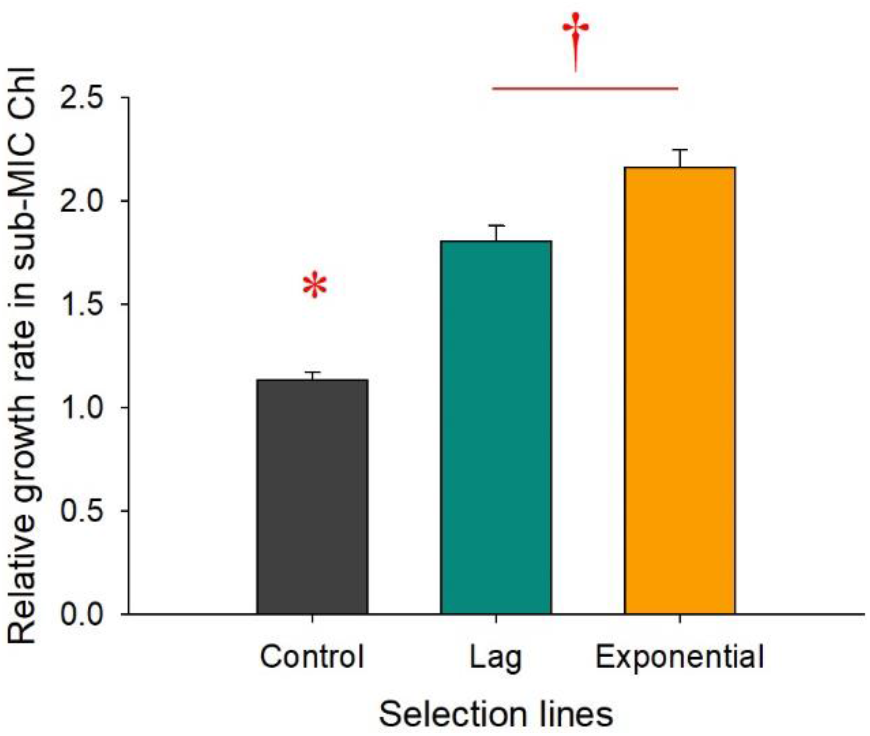
Relative growth rate of evolved populations in sub-MIC chloramphenicol. Bars represents mean of 12 values i.e., 6 replicate populations assayed twice. Whiskers represent ±SE. * denote control is significantly (p value < 0.05) less than lag and exponential in Tukey’s post-hoc analysis. † denotes significant differences between lag and exponential treatments in Tukey’s post-hoc analysis.

To summarize, the genomic changes due to selection for UV resistance did not result in fitness changes, between the treatments, in all but one novel environment. Interestingly, the genomic changes induced by selection under UV, never led to a decrease in fitness, w.r.t control populations, in the novel environments. In the case of chloramphenicol, where there was an observable phenotypic difference between the lag and the exponential treatments, the effect was not consistent as lag treatment had higher MIC whereas exponential treatment had higher growth rate. In other words, genomic differences between the populations that experienced selection in different growth phases did not translate into observable differences in their phenotypes, either in the selection environment (Figure 2), or in the novel environments (Figure 5).

## 4. Discussion

Bacterial growth phases show considerable variations in their resistance to ionizing radiations. While some studies have demonstrated that the exponential phase can have increased sensitivity to ionizing radiations (Morton & Haynes, 1969; Child *et al.*, 2002; Dantur & Pizarro, 2004; Bucheli-Witschel *et al.*, 2010; Arroyo *et al.*, 2012), others have shown that this phase can have greater resistance to such radiations (Keller & Maxcy, 1984; Hastings *et al.*, 1986; Kottemann *et al.*, 2005; DeVeaux *et al.*, 2007; Sukhi *et al.*, 2009). In this study, we compared the UV sensitivity of our ancestral strain of *Escherichia coli* MG1655 in lag and exponential phase. Ancestral cells in the exponential phase were more sensitive to UV, which resulted in larger reduction (~ 1-2 log_10_ fold more reduction) in colony forming units (CFUs) compared to lag phase (Fig. 1). The two growth phases differ in a number of factors that can influence UV sensitivity. Factors such as changes in growth rate (Keller & Maxcy, 1984; Berney *et al.*, 2006; Bucheli-Witschel *et al.*, 2010), growth environment (particularly nutrition availability post irradiation) (Child *et al.*, 2002; Sukhi *et al.*, 2009), and the quantity of genetic material (Fabre, 1970; Bucheli-Witschel *et al.*, 2010), have been suggested to explain the differential UV sensitivity in different growth phases. These differences can influence the organisms’ ability to repair UV induced damages and consequently the dynamics of resistance evolution.

We subjected our *E.coli* populations to selection under UV exposures during the lag and the exponential phase. After 100 rounds of exposure and growth, the UV exposed populations evolved increased resistance to UV compared to the non-exposed control populations (Fig. 2). 370 seconds of UV exposure in the ancestor and the control populations resulted in 5-7 log_10_reduction in viable CFUs whereas the UV-treated populations experienced only 1-2 log_10_reduction in CFUs (see supplementary material Table S7 for the raw data). Contrary to our expectations, the differences between growth phases observed in the sensitive strains (ancestor and control populations) did not translate into significant differences in the evolved response. Both the treatments had evolved resistance to UV in both growth phases, irrespective of the selection environment. These results are comparable with previous studies in *Bacillus subtilis* (Wassmann *et al.*, 2011) where it was shown that populations that evolved UV resistance in the stationary phase, also showed similar increased UV resistance in all growth phases. Interestingly, in that study, this was observed in spite of the fact that their ancestral and control populations showed growth phase dependent sensitivity to UV (Wassmann *et al.*, 2011). Considering the results from our study and Wassman et al., (Wassmann *et al.*, 2011), it is tempting to hypothesize that there are only a few routes to UV resistance, which might not be influenced by the growth phases. To investigate this possibility, we employed evolve and re-sequence (E&R) technique on replicate populations to characterize the genomics of UV resistance at different growth phases.

Both the treatments saw the accumulation of a large number of mutations with an overrepresentation of GC→AT transitions (at least 50% of all mutations). GC→AT transitions are the signature mutations of UV (Griffiths *et al.*, 2005; Brash, 2015) which are a result of the repair of UV-induced oxidative damages to the DNA (Wang *et al.*, 1998). This demonstrates the strong influence of UV radiation in shaping the genome of the UV-treated populations.

To investigate the genomic changes associated with UV resistance, we focused on the non-synonymous mutations in the populations. We found that a single amino acid change in *recA* (D161G) was convergent across all four UV-treated populations. Our ancestor had a mutation in *recA* (G161D) with respect to the reference strain, MG1655 (NC_000913.3). It is known that D161 is an extremely conserved amino acid and it plays an important role in determining the preference of recA for single stranded DNA over double stranded DNA (Shinohara *et al.*, 2015). The fact that all UV-treated populations, but not the controls, had fixed for reversion to the wild type form, suggests that this region is highly essential for the functioning of *recA* in the presence of UV stress.

While *recA* protein is a key regulator of recombinational repair of UV induced damages (Smith *et al.*, 1987) we did not find any other prominent change in its sequence. Interestingly, we found mutations in *recJ* and *recQ* genes, which are components of the recFOR recombination machinery that is regulated by *recA* in response to DNA damage (Morimatsu & Kowalczykowski, 2014). Additionally, mutations in the RNA polymerase (RNAP) genes, *rpoB* and *rpoC*, were also observed in UV-treated populations. Although mutation in rpoB gene are major effectors of rifampicin resistance, it was not observed in all populations. Since rifampicin resistance had evolved in both UV-treated populations, resistance cannot simply be explained by rpoB mutation. We also note that the mutation M1243L (in L3) is not one of the 69 known rpoB mutation conferring rifampicin resistance (Garibyan *et al.*, 2003). However, mutations in RNAP have previously been shown to confer radiation resistance in *Deinococcus* (Bruckbauer *et al.*, 2019). Studies show that RNAP and DNA repair proteins can interact when replication is stalled in rapidly dividing cells (Trautinger *et al.*, 2005; Baharoglu *et al.*, 2010). In our populations, we see an interesting combination of mutations in the recFOR pathway and RNAP genes. Mutations in recJ and rpoC were found together and at similar frequency in L1 and E3 populations while mutations in recQ and rpoB occurred together in L3, at similar frequency (see supplementary document S8). It is likely that the interaction between the different components of the two mechanisms (recFOR pathway and RNAP) is crucial for UV resistance. However, these components can interact in multiple ways and figuring out the details of these interactions is a challenge that is outside the scope of this study.

Although DNA/nucleotides are considered to be the primary target of UV radiation, there is growing evidence that cell membrane (Alper, 1977; Schwarz, 1998; Kumar *et al.*, 2016) and proteins (Krisko & Radman, 2010; Daly, 2012) are also indirect targets of UV damage via generation of reactive oxygen species (ROS). Prevention and/or tolerance to damages to the cell membrane and protein can be an alternate strategy of UV resistance. It has been shown that ROS generated by UVB stress causes lipid peroxidation resulting in cell membrane damage (Gomes *et al.*, 2013; Santos *et al.*, 2013). Thus, maintaining cell membrane integrity is probably one of the first priorities when under UV stress. For instance, in *Enterobacter cloacae*, outer membrane protein (*ompC*) and periplasmic oligopeptide binding protein (*oppA*) were among the differentially expressed genes when exposed to UVB (Kumar *et al.*, 2016). Although a causal link between cell membrane structure and radiation resistance has not been experimentally shown, the highly radio-resistant *Deinococcous* spp. is well known for its unique multilayered (six layers) cell membrane (Makarova *et al.*, 2001). Thus, it is possible that the structure and composition of the cell membrane might influence radiation resistance. Consistent with this, we see that all our UV-treated populations, but not the controls, carry mutations in *mepS*, an endopeptidase which is a part of cell wall biogenesis (Singh *et al.*, 2012). Additionally, mutation clusters in genes involved in or a part of cell membrane structure were consistent among UV-treated populations (Fig. 4). With the present data, it is not possible for us to comment on whether these mutations directly led to increased UV resistance or were neutrally accumulated as a consequence of increased mutation rate in the UV-treated populations. However, this opens potential avenues for investigating the mechanism of UV resistance from the point of view of cell membrane structure and composition. It is also possible that the increased mutation supply in the UV treated populations allowed them to explore the mutation landscape for mutations that could be beneficial in the laboratory growth conditions. This is evident from the fact that both UV-treated populations had significant increase in growth rate in nutrient broth than the control populations (Fig. 5A).

One of the mechanisms of radiation resistance in *Deinococcus* is the export of damaged/degraded DNA following radiation (Battista, 1997; White *et al.*, 1999). Transport of the damaged oligonucleotides can prevent them from being reincorporated during repair. While the cells export damaged DNA, nutrients such as amino acids, sugars and phosphates may also be imported into the cell (Makarova *et al.*, 2001). This increased nutrition is essential for the energy expensive DNA repair process (Venkateswaran *et al.*, 2000). Consequently, radio-resistance has also been attributed to efficient transport of nutrients into the cell (Makarova *et al.*, 2001; Child *et al.*, 2002; Sukhi *et al.*, 2009). In line with this, we observe clustering of mutations in genes involved in cellular transport (Fig. 4). These mutations are unique to the lag treatment and were not present in the exponential treatment. Another unique characteristic of the genome of the lag treatment is the fixation of mutations in *lexA* genes. *lexA* is a transcriptional repressor of SOS response (Maslowska *et al.*, 2019). Additionally, mutations in transcriptional regulators such as *crp*, *deoR*, *fadR*, *hfq* were also common in the lag populations (Table 2). Taken together, this suggests that in addition to the direct response to radiation, UV resistance in lag treatment can comprise of protection/tolerance mechanisms as well as indirect response via regulation of other genes.

The mutations in the exponential treatment, except for those in genes related to repair and cell membrane structure and function, could not be associated with any known pathway associated with UV resistance (Table 2 and Fig. 4). Characterizing the role of signal transduction, cell adhesion, metal-ion and nucleotide binding proteins, and enzymes such as kinases, and serine esterase, in the context of UV resistance, might suggest the association of novel pathways of UV resistance.

Horizontal gene transfer (HGT), by natural transformation of exogeneous DNA from the environment, is known to be induced by stress (Claverys *et al.*, 2006; Prudhomme *et al.*, 2006), particularly UV radiation (Charpentier *et al.*, 2011). Additionally, bacterial competence is also known to be influence by multiple factors including, growth phase (Szostkova *et al.*, 1999). However, exploring the influence of HGT on the genome evolution of our UV-treated populations was out of the scope of this study.

UV as a mutagen is expected to increase the genome wide mutagenesis. A large number of the resulting mutations are expected to be deleterious in the selection environment and therefore likely to be purged by purifying selection. The observed dN/dS ratio of mutations in the coding region being less than one suggests that the UV-treated populations were indeed subjected to purifying selection (Table 1). The mutations that escaped being purged and accumulated to high frequencies were either beneficial or neutral in the selection environment. However, it is possible that some of these neutral mutations are contextually neutral, i.e. have an effect on fitness when the environment is altered (Wagner, 2005). To investigate this possibility, we studied the fitness of the UV-treated populations under various antibiotics, heavy metals and carbon sources.

Despite the large genetic variation, fitness of the UV-treated populations were significantly different from the control populations in only four out of 12 novel environments namely, chloramphenicol, nalidixic acid, rifampicin and cobalt chloride (Figure 5). One possible way by which the UV-treated populations could have acquired resistance to these environments is via the UV induced alterations in the cell membrane permeability. Outer membrane permeability has previously been implicated in the evolution of antibiotic resistance (Delcour, 2009; Ghai & Ghai, 2018; May & Grabowicz, 2018). On the other hand, exposure to UV could have also resulted in the introduction of antibiotic resistance mutations in smaller subpopulations which could allow them to grow at higher concentrations of antibiotics (Band & Weiss, 2019). Interestingly, the lag and exponential populations showed significant differences only in resistance to chloramphenicol. Even in terms of chloramphenicol resistance, the differences between lag and exponential treatments were not consistent across MIC and growth rate (Figures 5 and 6). Thus, taken together, the genomic signatures of UV adaptation in lag and exponential treatment populations did not result in any major phenotypic differences between them in both the selection as well as novel environments.

## 5. Conclusion

Experimental evolution in combination with high-throughput sequencing (evolve and resequence) is an extremely powerful tool to study the genomics of adaptation (Long *et al.*, 2015; Schlötterer *et al.*, 2015). While it is known that the phenotype-genotype map can be degenerate, molecular parallelisms can be found at different levels of genome organization ranging from same nucleotide substitution to similar gene networks (Rosenblum *et al.*, 2014; Hao *et al.*, 2019). Evidence for this comes from previous studies where huge diversity in the beneficial mutations have been reported at the level of nucleotides but convergence was observed at the level of genes and functional groups (Woods *et al.*, 2006; Tenaillon *et al.*, 2012). Similarly, in our study, although the large number of mutations initially seemed to be randomly distributed in the genome, we observed some convergence of functional groups. DNA repair, RNA polymerase and genes associated with cell membrane structure were some of the convergent changes observed in the UV-treated populations. However, the exposure to UV during different growth phases also led to some unique genomic signatures. It is likely that the two growth phases represent different physiological and biochemical environments inside the cell, which could have constrained the UV induced mutations and consequently the amount and nature of genetic variation available for selection to act. Nevertheless, mutations in genes for mechanisms besides DNA repair systems that have been observed in the UV-treated populations are suggestive of the role of other indirect pathways involved in UV resistance. These results are reminiscent of mechanisms of extreme radio-resistance in *Deinococcus radiodurans* R1 where resistance has been shown to rely more on indirect mechanisms such as cellular cleansing, signal transduction and transcriptional regulation than on extensive damage repair mechanisms (Makarova *et al.*, 2001; Galperin *et al.*, 2006; Blasius *et al.*, 2008). In addition to recognizing the different possible mechanisms of UV resistance, this study highlights the influence of physiology in shaping the genomic evolution. We see that mutational profiles are dependent on the growth phase of exposure. Very little is known about such growth phase specific effect of most mutagens. Such physiological biases of mutagenesis can have important implications in industrial strain improvement studies. Additionally, as UV radiation is widely used as a disinfectant, it is important to acknowledge the evolution of antibiotic resistance as a correlated response. To better manage this, further experiments need to be done to understand the relationship between growth physiology, UV response and antibiotic resistance evolution.

## Acknowledgements

We thank Gayathri Pananghat, Nishad Matange, Krishanpal Karmodiya, Farhat Habib, and Madhusudhan M. S. for their valuable inputs. SS was supported by a Junior Research Fellowship initially sponsored by IISER Pune and then by Department of Biotechnology (DBT), Govt. of India. KL was supported by a KVPY fellowship, sponsored by the Department of Science and Technology (DST), Govt. of India. This project was supported by a grant from Department of Biotechnology, Government of India (#BT/PR22328/BRB/10/1569/2016) and internal funding from IISER Pune. The authors declare no conflict of interest.

## Supplementary material

**Figure S1.**
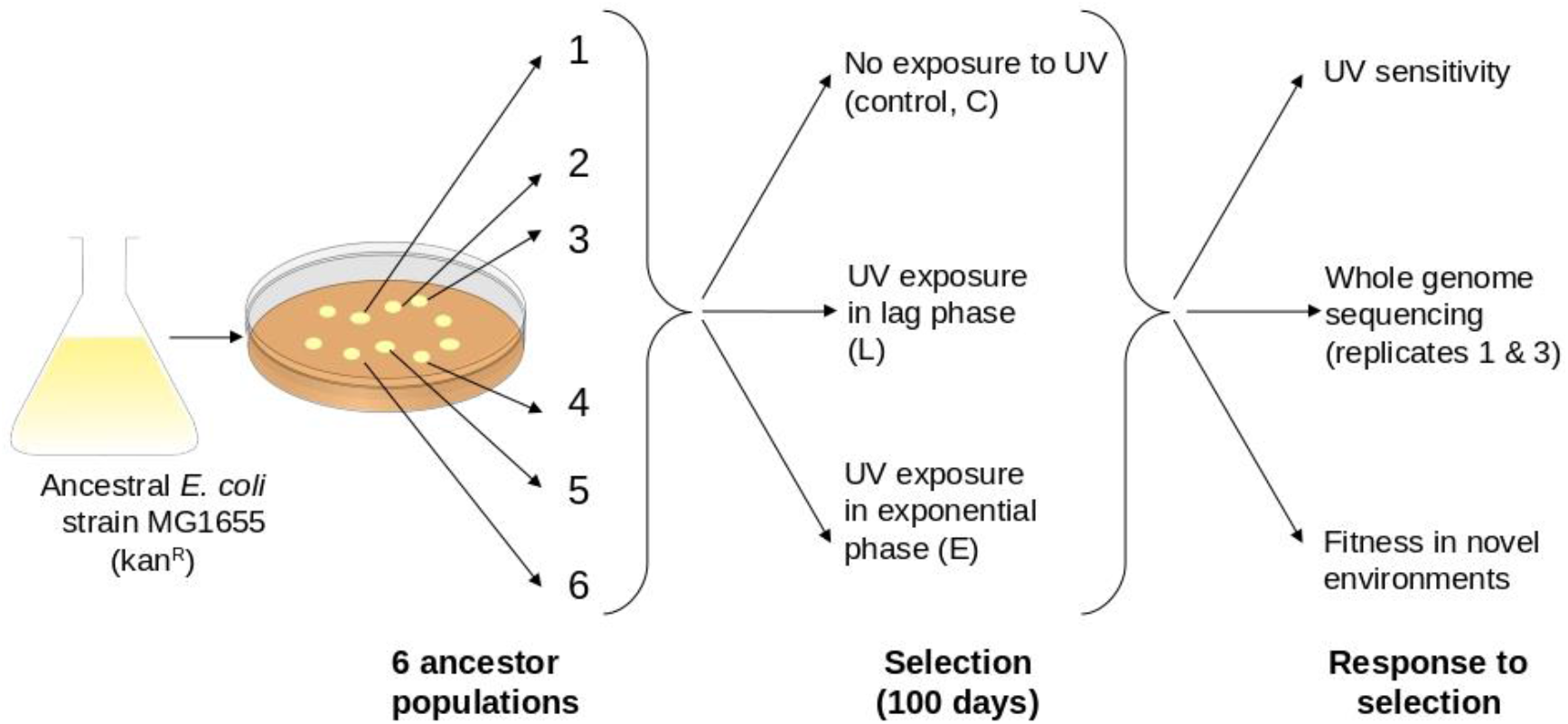
An overview of the experimental design in this study.

**Table S2.** Exposure duration during selection. The duration of exposure to UV was increased every 5 days.

| **Days** | **UV exposure duration (seconds)** |
| --- | --- |
| 1 - 5 | 15 |
| 6 - 10 | 20 |
| 11 - 15 | 25 |
| 16 - 20 | 35 |
| 21 - 25 | 45 |
| 26 - 30 | 60 |
| 31 - 35 | 75 |
| 36 - 40 | 90 |
| 41 - 45 | 110 |
| 46 - 50 | 130 |
| 51 - 55 | 160 |
| 56 - 60 | 200 |
| 61 - 65 | 220 |
| 66 - 70 | 240 |
| 71 - 75 | 270 |
| 76 - 80 | 300 |
| 81 - 85 | 330 |
| 86 - 90 | 340 |
| 91 - 95 | 350 |
| 96 - 100 | 370 |

**Table S3.**
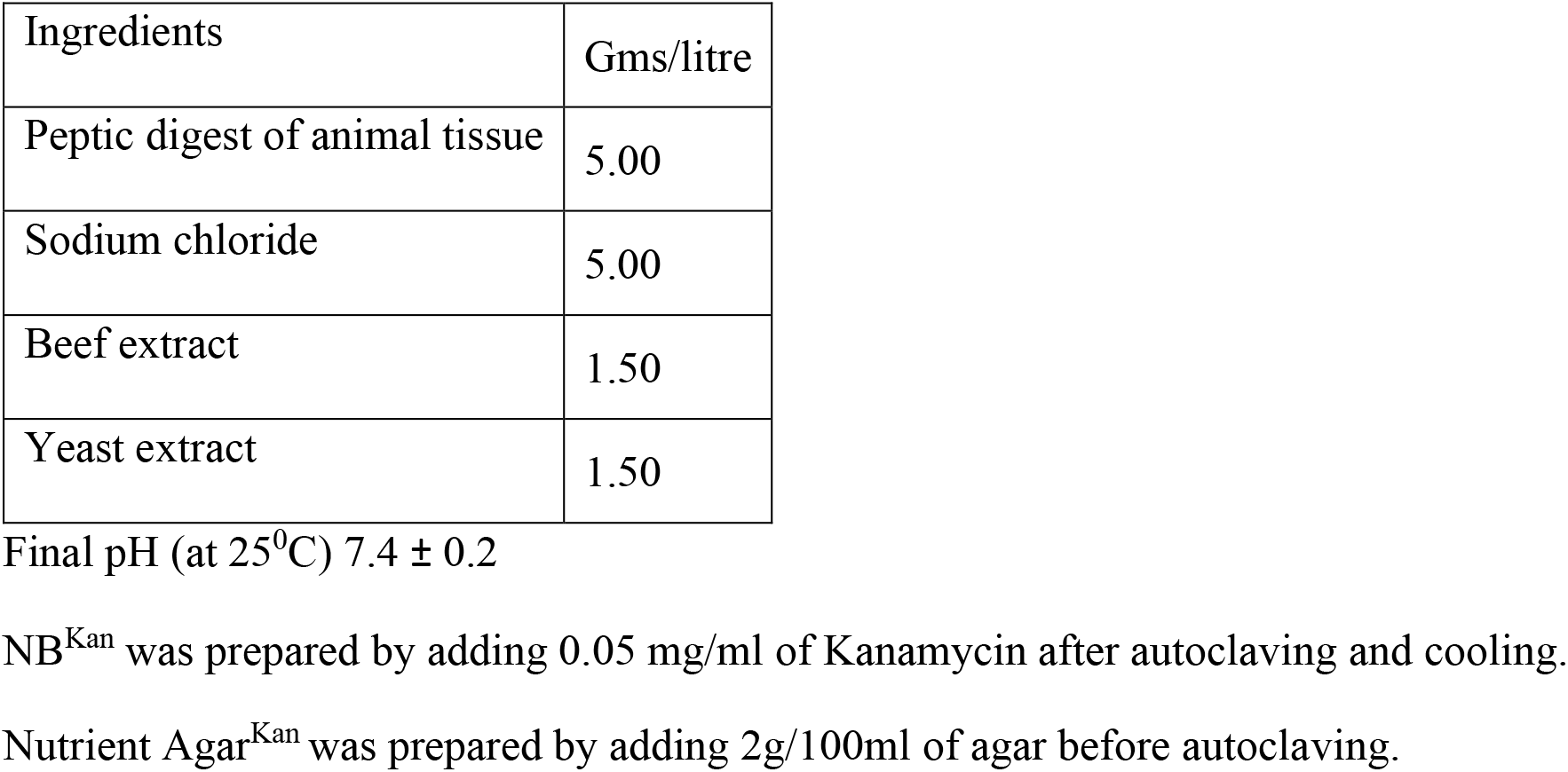
Composition of Nutrient broth.

**Table S4.** Composition of M9 media. The growth rate of the selected populations were measured in five minimal media (M9) with fructose, glycose, glycerol, mannose and thymine as single carbon source.

| Ingredients | Gms/litre |
| --- | --- |
| NaH<sub>2</sub>PO<sub>4</sub>.7H<sub>2</sub>O | 12.8 |
| KH<sub>2</sub>PO<sub>4</sub> | 3.0 |
| NaCl | 0.5 |
| NH<sub>4</sub>Cl | 1.0 |
| MgSO<sub>4</sub> | 0.24 |
| CaCl<sub>2</sub> | 0.01 |
| Carbon source | 4 |

0.05 mg/ml of Kanamycin was added after autoclaving and cooling.

**Table S5.**
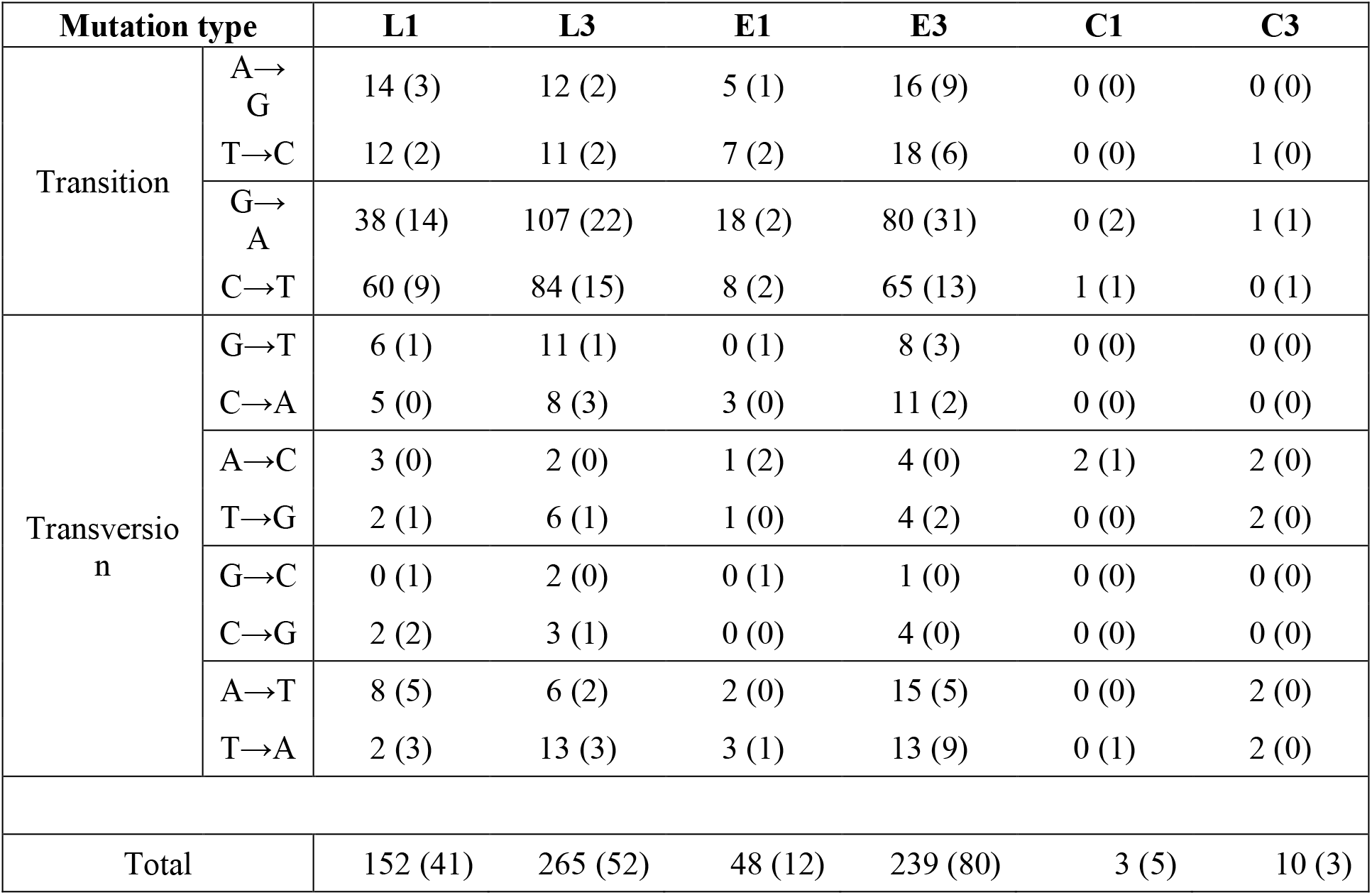
Mutational spectrum of all SNPs. The mutational spectrum of all SNPs, including synonymous and non-synonymous mutations, identified in the selected populations in the coding regions (intergenic regions). The spectrum of two replicates (1 and 3) of lag (L), exponential (E) and control (C) populations are listed below.

**Table S6.**
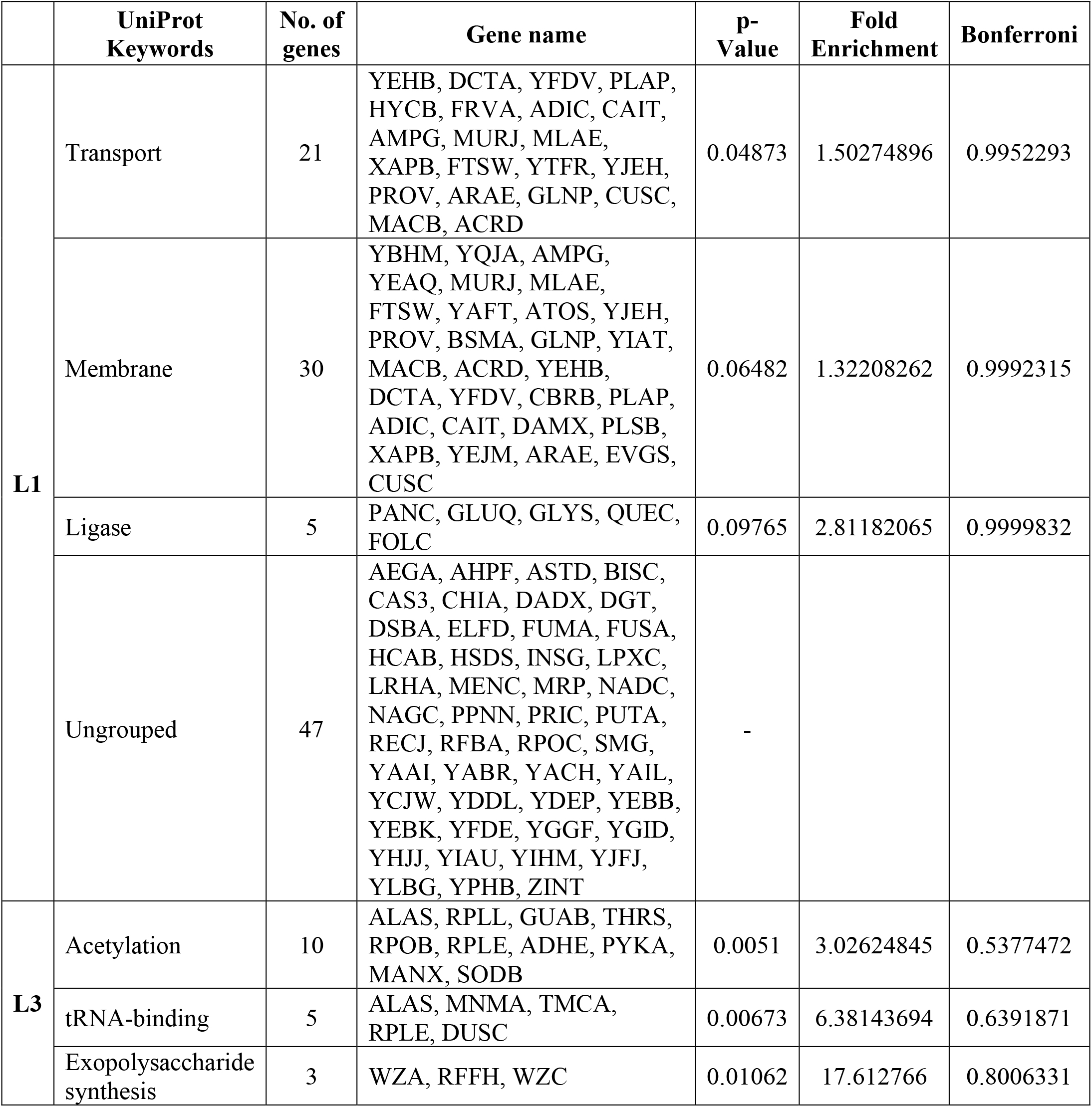
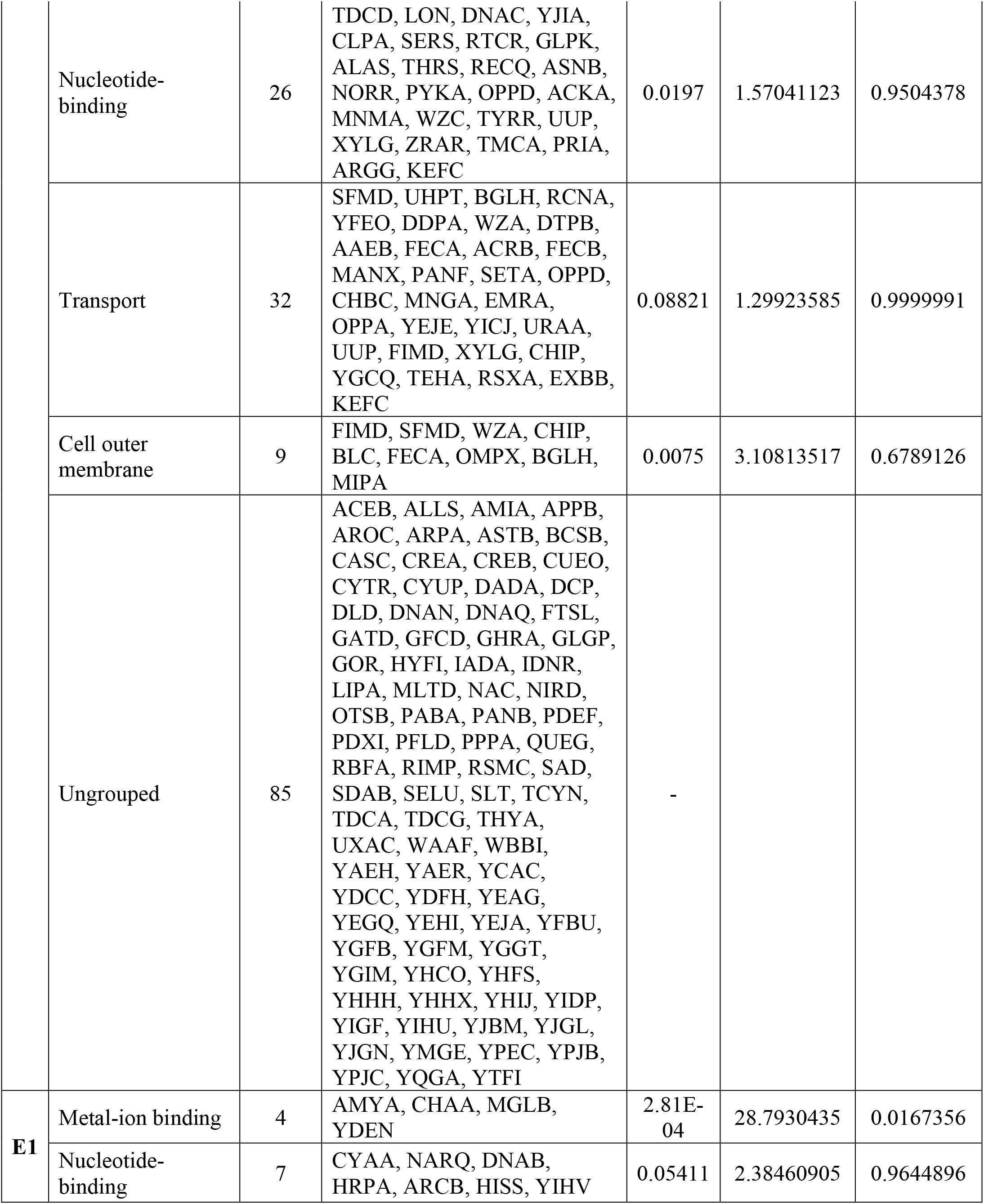
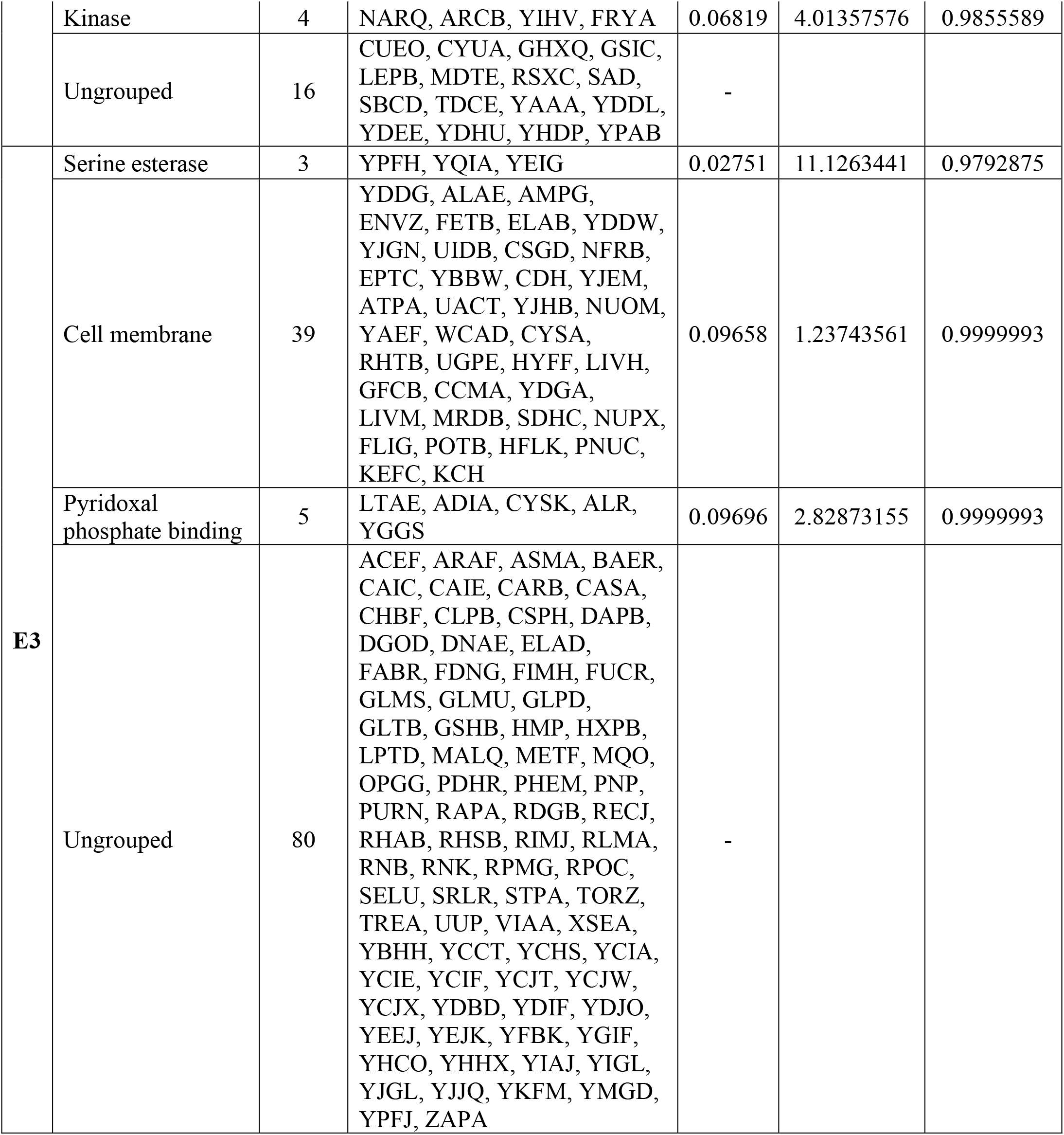
Functional categorization and enrichment analysis using DAVID. The list of genes that were unique to each treatment population categorized into functional groups using DAVID. The enrichment analysis did not yield significant results as seen from the fold enrichment and Bonferroni p-values (corrected for inflated family-wise error rates). This could be a result of the high proportion (~50%) of ungrouped genes.

**Table S7.** UV sensitivity values. UV sensitivity of the evolved, control and ancestral population when exposed to 370 seconds of UV.

| **Trial** | **Assay-Growth phase** | **Selection-Growth phase** | **Replicate** | **UV induced $\log_{10}$ reduction** |
| --- | --- | --- | --- | --- |
| I | Lag | Lag | 1 | 0.836497868 |
| I | Lag | Lag | 2 | 0.954471146 |
| I | Lag | Lag | 3 | 0.726998728 |
| I | Lag | Lag | 4 | 0.466263327 |
| I | Lag | Lag | 5 | 1.093049592 |
| I | Lag | Lag | 6 | 0.434410528 |
| II | Lag | Lag | 1 | 1.029383778 |
| II | Lag | Lag | 2 | 1.761117911 |
| II | Lag | Lag | 3 | 0.645838916 |
| II | Lag | Lag | 4 | 0.00152652 |
| II | Lag | Lag | 5 | 1.79278894 |
| II | Lag | Lag | 6 | 2.407156813 |
| I | Lag | Exponential | 1 | 1.698970004 |
| I | Lag | Exponential | 2 | 0.812326868 |
| I | Lag | Exponential | 3 | 1.268306151 |
| I | Lag | Exponential | 4 | 1.597943425 |
| I | Lag | Exponential | 5 | 1.12082217 |
| I | Lag | Exponential | 6 | 1.660812302 |
| II | Lag | Exponential | 1 | 2 |
| II | Lag | Exponential | 2 | 1.643922438 |
| II | Lag | Exponential | 3 | 0.672347524 |
| II | Lag | Exponential | 4 | 1.191282758 |
| II | Lag | Exponential | 5 | 1.818858748 |
| II | Lag | Exponential | 6 | 2.483281563 |
| I | Lag | Control | 1 | 4.789146635 |
| I | Lag | Control | 2 | 6 |
| I | Lag | Control | 3 | 4.558490387 |
| I | Lag | Control | 4 | 4.853476972 |
| I | Lag | Control | 5 | 5.875061263 |
| I | Lag | Control | 6 | 4.962211439 |
| II | Lag | Control | 1 | 4.768839833 |
| II | Lag | Control | 2 | 4.788455999 |
| II | Lag | Control | 3 | 5.67516709 |
| II | Lag | Control | 4 | 5.818156412 |
| II | Lag | Control | 5 | 7.243038049 |
| II | Lag | Control | 6 | 4.953895213 |
| I | Lag | Ancestor | 1 | 5.736758565 |
| I | Lag | Ancestor | 2 | 5 |
| I | Lag | Ancestor | 3 | 4.911273436 |
| I | Lag | Ancestor | 4 | 5.780857144 |
| I | Lag | Ancestor | 5 | 5.795880017 |
| I | Lag | Ancestor | 6 | 4.848531216 |
| II | Lag | Ancestor | 1 | 5.836745966 |
| II | Lag | Ancestor | 2 | 5.716003344 |
| II | Lag | Ancestor | 3 | 5.25224605 |
| II | Lag | Ancestor | 4 | 6.176091259 |
| II | Lag | Ancestor | 5 | 7.041392685 |
| II | Lag | Ancestor | 6 | 5.281846679 |
| I | Exponential | Lag | 1 | 2.050609993 |
| I | Exponential | Lag | 2 | 2.965237894 |
| I | Exponential | Lag | 3 | 1.437750563 |
| I | Exponential | Lag | 4 | 1.852323676 |
| I | Exponential | Lag | 5 | 2.30207272 |
| I | Exponential | Lag | 6 | 1.446200644 |
| II | Exponential | Lag | 1 | 2.105633581 |
| II | Exponential | Lag | 2 | 2.49574251 |
| II | Exponential | Lag | 3 | 1.635261445 |
| II | Exponential | Lag | 4 | 1.897804391 |
| II | Exponential | Lag | 5 | 2.416489212 |
| II | Exponential | Lag | 6 | 1.249877473 |
| I | Exponential | Exponential | 1 | 2 |
| I | Exponential | Exponential | 2 | 2.486849858 |
| I | Exponential | Exponential | 3 | 0.654996067 |
| I | Exponential | Exponential | 4 | 2.266052412 |
| I | Exponential | Exponential | 5 | 1.67593102 |
| I | Exponential | Exponential | 6 | 2.006182232 |
| II | Exponential | Exponential | 1 | 2.817581758 |
| II | Exponential | Exponential | 2 | 4.628057027 |
| II | Exponential | Exponential | 3 | 1.137926523 |
| II | Exponential | Exponential | 4 | 3.696264111 |
| II | Exponential | Exponential | 5 | 1.879979713 |
| II | Exponential | Exponential | 6 | 2.241153905 |
| I | Exponential | Control | 1 | 5.642488648 |
| I | Exponential | Control | 2 | 7.301029996 |
| I | Exponential | Control | 3 | 7.694605199 |
| I | Exponential | Control | 4 | 7.079181246 |
| I | Exponential | Control | 5 | 7.08754481 |
| I | Exponential | Control | 6 | 7.63973544 |
| II | Exponential | Control | 1 | 6.970897618 |
| II | Exponential | Control | 2 | 6.388248237 |
| II | Exponential | Control | 3 | 7.630427875 |
| II | Exponential | Control | 4 | 6.621365147 |
| II | Exponential | Control | 5 | 6.722908012 |
| II | Exponential | Control | 6 | 6.5633732 |
| I | Exponential | Ancestor | 1 | 8.335792102 |
| I | Exponential | Ancestor | 2 | 8.432969291 |
| I | Exponential | Ancestor | 3 | 8.359835482 |
| I | Exponential | Ancestor | 4 | 7.698970004 |
| I | Exponential | Ancestor | 5 | 6.638272164 |
| I | Exponential | Ancestor | 6 | 8.170261715 |
| II | Exponential | Ancestor | 1 | 6.133893579 |
| II | Exponential | Ancestor | 2 | 7.996730515 |
| II | Exponential | Ancestor | 3 | 8.404833717 |
| II | Exponential | Ancestor | 4 | 8.118099312 |
| II | Exponential | Ancestor | 5 | 8.522878745 |
| II | Exponential | Ancestor | 6 | 7.750122527 |

**Figure S8.**
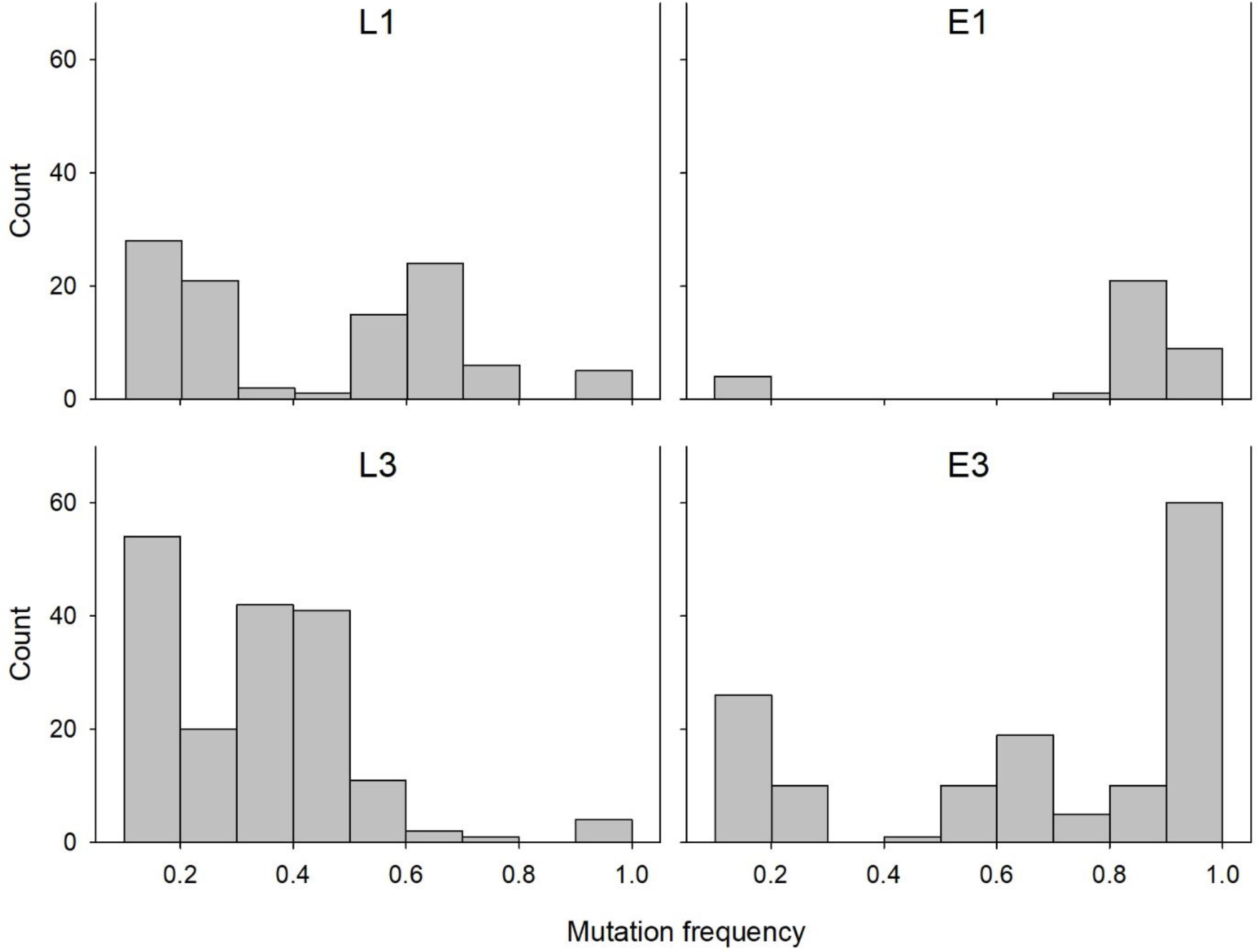
Distribution of mutation frequency in the UV selected populations. A histogram showing the distribution of the mutation frequency in the four UV-treated populations.

## Notes

### Competing Interest Statement

The authors have declared no competing interest.

